# Genetic disruption of isocitrate dehydrogenase arrests the full development of sexual stage parasites in *Plasmodium falciparum*

**DOI:** 10.1101/2022.08.05.502965

**Authors:** Ming Yang, Hangjun Ke, Joanne M. Morrisey, Michael W. Mather, Akhil B. Vaidya

## Abstract

The malaria parasites encode all eight mitochondrial enzymes that can constitute the classical tricarboxylic acid (TCA) cycle. We previously reported that genetic ablation of six out of eight TCA enzymes had no significant impact on the parasite survival and growth in asexual blood stage; aconitase knockout (*ΔACO*) parasites were, however, arrested as late-stage gametocytes. Herein, we describe a defective gametocyte development phenotype resulting from the knockout of another TCA enzyme, isocitrate dehydrogenase (IDH). Similar to the *ΔACO* gametocytes, *ΔIDH* gametocytes stall at stage III/IV and cannot form mature gametocytes or undergo exflagellation. Both *ACO* and *IDH* KO gametocytes exhibit fragmented mitochondrial morphology. Knockout of *IDH* also exerts a fitness cost, leading to a decrease in growth rate in erythrocytic stages. We examined two possible causes for the phenotype in the *ΔIDH* gametocytes: 1. Accumulation of reactive oxygen species (ROS) within the parasite mitochondrion due to diminished production of mitochondrial NADPH during the 10-14 days of gametocyte development; or 2. Cytotoxicity of built-up citrate upstream of IDH. We found that antioxidant agents such as N-acetylcysteine and MitoQ did not rescue the phenotype of *ΔACO* or *ΔIDH* gametocytes. Large amounts of citrate in the medium also had no significant effect on the gametocyte development or gamete formation of wildtype parasites. Our results demonstrate the importance of IDH in gametocytogenesis, providing evidence that IDH as well as aconitase could be further explored as targets for gametocidal agents and malaria transmission blockers.

## Introduction

Malaria remains one of the leading causes of morbidity and mortality worldwide [1], leading to estimated 244 million clinical cases and 627,000 deaths in 2020, a 12% increase in mortality over the previous year [2]. Despite global efforts to control and eradicate malaria, the continuous emergence of drug-resistant parasite strains [3] and insecticide resistant mosquitoes [4], as well as lack of an effective vaccine, underscore the need for novel antimalarial drugs and transmission blockers. *Plasmodium falciparum* causes the most severe form of malaria in humans. The asexual cyclic replication of the parasites within erythrocytes leads to the clinical symptoms of the disease, while gametocytes, the sexual stages of the parasites, are responsible for infecting a female *Anopheles* mosquito, in which the parasites undergo sexual recombination and insect-stage development to form sporozoites that in turn infect humans to perpetuate the life cycle. Most antimalarial drugs against asexual erythrocytic parasites have little or no effect on gametocyte development, allowing continued malaria transmission to occur after the clearance of asexual parasites [5]. Some drugs, such as chloroquine and quinine, kill early stage gametocytes through their action to disrupt the detoxification of free heme from hemoglobin digestion, but have no effect on later stages [6]. Similarly, atovaquone, an inhibitor of the cytochrome *bc*_*1*_ complex of the parasite mitochondrial electron transport chain (mtETC), acts against early but not mature gametocytes [7]. In spite of the rapid action of artemisinin and its ability to reduce gametocyte carriage [8, 9], transmission can still occur following artemisinin-based combination therapy [10]. Primaquine kills mature gametocytes [11-13] and was widely used as a single-dose drug to reduce malaria transmission prior to the introduction of artemisinin. However, primaquine is associated with a major safety issue in that it can cause severe hemolysis in patients with glucose-6-phosphate dehydrogenase deficiency [13]. With a growing recognition of the importance of controlling malaria transmission [14], identification of essential genes for gametocyte development will enable the screening for new compounds against developing and mature gametocytes.

The mitochondrion of *Plasmodium* parasites is an essential organelle for parasite survival and is highly divergent from the human counterpart [15]. Within the parasite mitochondrion, there are two validated antimalarial drug targets, the mtETC and the pyrimidine biosynthesis enzyme dihydroorotate dehydrogenase (DHODH). There are various chemical classes of inhibitors that disrupt mtETC function and collapse the mitochondrial membrane potential, including the clinical antimalarial atovaquone [16, 17] and a pre-clinical candidate compound ELQ-300 [18]. DHODH inhibitors such as DSM-265 have also undergone clinical trials [19]. While mitochondrial functions have been successfully targeted in asexual blood-stage parasites, the role of the mitochondrion during gametocyte development has not been fully characterized. The mitochondrion in *P. falciparum* gametocytes and ookinetes contains a higher density of tubular cristae than that of asexual parasites [20, 21], suggesting a possible shift to greater reliance on mitochondrial functions during gametocyte development and subsequent mosquito stages. This is also supported by the observation that the gametocyte mitochondrion drastically expands in size and forms a large branching structure as the gametocyte matures through stages I-V [22]. Furthermore, a study of *P. falciparum* gametocyte transcriptomes revealed that mRNAs of the mitochondrial TCA cycle enzymes are present in gametocytes [23] and proteomics data showed accumulation of TCA enzymes in mature gametocytes [24].

Previously, we reported that genetic ablation of six out of eight TCA enzymes did not have a significant effect on the asexual blood-stage parasites, but the TCA cycle was essential for the mosquito stages [25]. Interestingly, knockout of aconitase resulted in impaired gametocyte development [25]. Aconitase knockout gametocytes arrested at stage III and failed to mature into stage V [25]. This is consistent with the observation that chemical inhibition of parasite aconitase by sodium fluoroacetate (NaFAc) prevents gametocyte maturation [26]. Surprisingly, knockout of another TCA enzyme, α-ketoglutarate dehydrogenase (KDH), had no effect on sexual development [25], suggesting that aconitase activity by itself, rather than the complete turning of the TCA cycle, is critical during gametocytogenesis. Aconitase converts citrate to isocitrate. IDH then converts isocitrate to α-ketoglutarate, generating reduced nicotinamide adenine dinucleotide phosphate (NADPH). Unlike many other organisms, including humans, malaria parasites encode only one IDH enzyme (PF3D7_1345700), shown to be mitochondrially targeted and NADP^+^-dependent [27]. Thus, IDH is the only known source of NADPH within the parasite mitochondrion. NADPH is a major electron donor for glutathione, which defends against oxidative damage, thus, playing a crucial role in the maintenance of redox balance [28]. NADPH also serves as an important cofactor for NADPH-dependent enzymes that are involved in crucial biosynthetic processes. On the other hand, ablation of ACO also leads to citrate accumulation. Thus, the defects in *ΔACO* gametocytes could be due to the lack of mitochondrial NADPH production or the accumulation of potentially cytotoxic citrate. Furthermore, ACO is known to play extra-mitochondrial roles in mammals through translational control of mRNAs bearing iron-regulatory elements [29], and if playing a similar role in *Plasmodium*, its absence might affect the gametocyte development program.

To further explore the role of TCA enzymes in gametocyte development, we utilized genetic approaches to study the NADPH-generating isocitrate dehydrogenase. We previously knocked out *IDH* in the D10 line of *P. falciparum* [25], a strain that does not form gametocytes; thus, we were unable to study the role of IDH in the gametocyte stage. Here, we report the deletion of the *IDH* gene in the gametocyte-competent strain of *P. falciparum*, NF54, to help assess the role of IDH in gametocytogenesis. In the asexual blood stage, disruption of *IDH* exacted a fitness cost on NF54 parasite growth. More importantly, *ΔIDH* gametocytes stalled at stage III and could not mature into stage V or form gametes, suggesting that IDH is essential for gametocyte development. We then pursued experiments designed to shed light on the possible cause(s) of compromised gametocytogenesis in *ΔACO* and *ΔIDH* gametocytes.

## Materials and methods

### Plasmid construction

The strategy used for *IDH* (PF3D7_1345700) gene knockout by CRISPR-Cas9-induced double crossover homologous recombination is illustrated in S1 Fig panel A. A 5’ homologous region (HR) and a 3’ HR that flank the segment of the *IDH* gene to be deleted were cloned [25] into the pCC1 plasmid [30], which contains human dihydrofolate reductase (*hDHFR*) as a selectable marker that confers resistance to WR99210 (5nM), an inhibitor of the parasite DHFR [31]. A list of prospective 20 base sequences (N_20_) for guide RNAs was generated using the Eukaryotic Pathogen CRISPR guide RNA (gRNA) design tool (http://grna.ctegd.uga.edu/) that target the *IDH* chromosomal DNA segment flanked by the 5’ and 3’ HRs. The three independent N_20_ sequences with highest total scores (S1 Table: PfIDHgRNA1_N20, PfIDHgRNA2_N20, PfIDHgRNA3_N20) were individually cloned between the BtgZI sites in pAIO, which carries both a gRNA expression cassette and a yeast dihydroorotate dehydrogenase-Cas9 endonuclease expression cassette [32]. Primers used for plasmid construction are shown in S1 Table. Cloned DNA inserts were verified by DNA sequencing.

### Parasite lines, parasite culture and transfection methods

*P. falciparum* NF54 was the parental line used to generate *ΔIDH* parasites. The *ΔACO* NF54 line was previously reported [25]. Parasites were cultured in human O^+^ erythrocytes at 5% hematocrit in RPMI 1640 medium supplemented with 0.5% AlbuMAX^®^ and incubated at 37 °C under low oxygen conditions (89% N_2_, 5% CO_2_ and 6% O_2_). Human O^+^ erythrocytes were purchased from Interstate Blood Bank, Inc. (Memphis, TN). Transfections were carried out using the standard method [33]. Briefly, ring-stage parasites from a culture at ≈5% parasitemia were washed three times with pre-warmed Cytomix (pH=7.6) and then resuspended with an equal volume of ice-cold Cytomix. An aliquot of 250 μl parasite suspension was mixed with 50 μg of each of the pAIO-derived plasmids containing one of three independent *IDH*-specific gRNA sequences and 12 μg linearized pCC1-PfIDH plasmid (digested by HincII, see Figure 1A) in a 0.2 cm cuvette. Electroporation was carried out in a Bio-Rad Genepulser set at 0.31 kV and 950 μF. Parasite cultures were exposed to 5 nM WR99210 starting 48-hour post-electroporation and split weekly until viable parasites were observed by Giemsa-stained blood smears.

**Fig 1.**
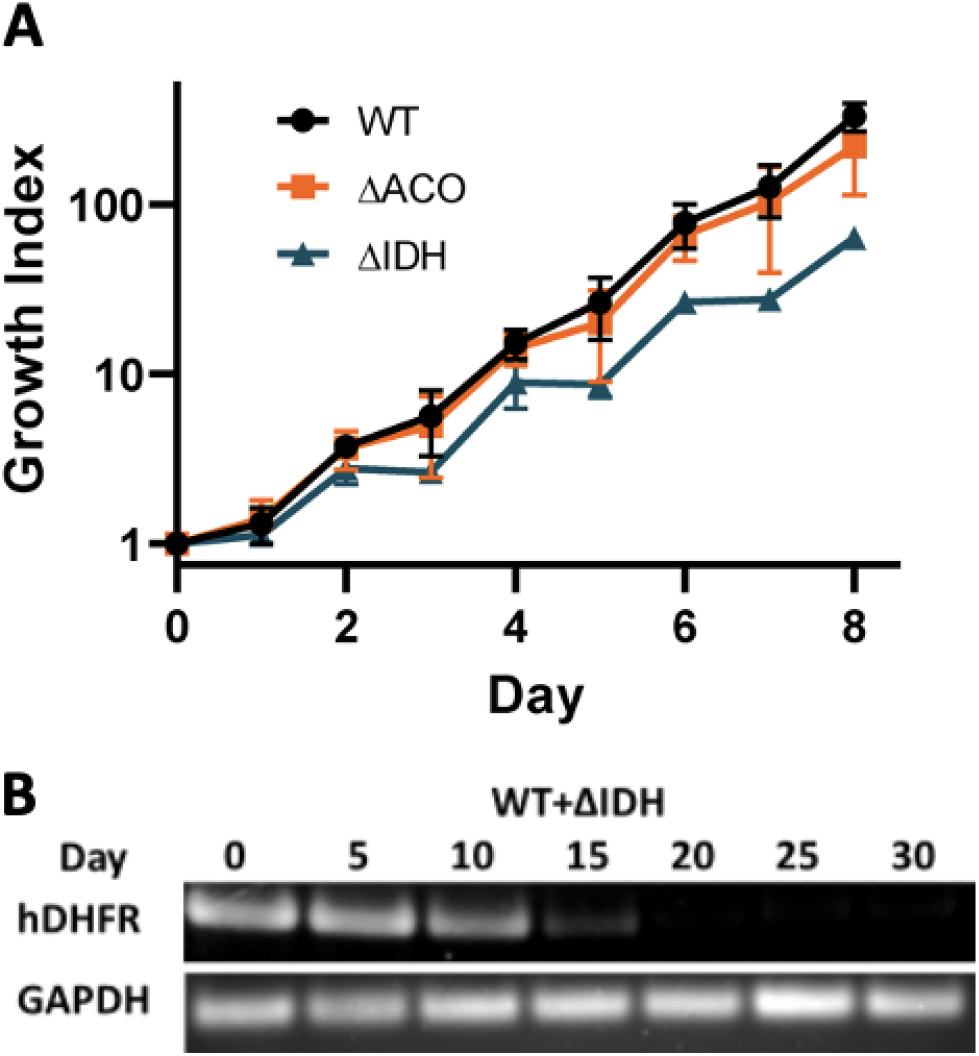
Phenotypic characterization of *ΔIDH* in the asexual blood stage. (A) Growth curves of NF54 wildtype, *ΔACO* and *ΔIDH* asexual blood stage parasites (mean and SEM plotted at each time point from 2 biological replicates of each line). (B) Diagnostic PCR assessing the PCR products of *hDHFR* and *PfGAPDH* genes in the fitness cost assay. WT and *ΔIDH* asexual blood-stage parasites were mixed at a 1:1 ratio on day 0, and samples are collected every 5 days at day 0, 5, 10, 15, 20, 25 and 30 for isolation of genomic DNA. Primers used are shown in S1 Table. Full agarose DNA gel images are shown in S2 Fig.

### DNA isolation and PCR

Parasite genomic DNA was isolated from trophozoite-stage parasites using the QIAamp DNA blood Mini kit (Qiagen). For PCR, each reaction was set up in a 25 μl volume containing 2.5 μl 10X reaction buffer, 1.5 μl forward primer (5 μM), 1.5 μl reverse primer (5 μM), 1 μl 10 mM dNTPs, 0.3 μl Taq polymerase, 100 ng genomic DNA and sterilized deionized water. The thermal cycling program was set as follows: 2 min denaturation at 95 °C, 30 cycles of 30 sec denaturation at 95 °C, 45 sec annealing at 50 °C and 5 min extension at 65 °C, and a final 10 min extension at 65 °C. Pfu Turbo DNA polymerase (Agilent Technologies) was used to amplify parasite DNA for cloning. PCR primers are shown in S1 Table.

### Growth curves

NF54 WT and knockout parasites (*ΔACO* and *ΔIDH*) were tightly synchronized using 0.5 M Alanine/10 mM HEPES (pH 7.5) for multiple rounds and seeded at 1% ring-stage parasites on day 1. From day 2 to day 9, parasites were collected daily and fixed overnight at 4 °C with 4% formaldehyde in phosphate buffered saline (PBS). All fixed parasite samples were stained with SYBR® Green I Nucleic Acid Gel Stain (ThermoFisher Scientific) at 1:1,000 dilutions for 2 hours at room temperature and then washed three times with PBS. 1 μl of the stained parasites pellet was added into 1 ml of sterilized deionized water and the samples were run through the BD Accuri™ C6 Plus flow cytometer. The sample acquisition was set as follows: run with a limit of 1,000,000 events with ungated samples; fluidics speed was set at medium; a threshold of 18,000 was set on FSC-H for all samples; cell density during acquisition was adjusted to approximately 5,000 events/μl. The parasitemia was measured as the percentage of SYBR-positive cells in 1 million RBCs indicated by distinct peaks with mean fluorescent intensity greater than 10^5^ (R.U.). On Days 4, 6 and 8, cultures were spilt 1:5. Growth index (Y-axis) was calculated by multiplying the parasitemia by the splitting factor 5.

### Fitness cost assay

Tightly synchronized trophozoite-stage NF54 WT parasites (≈5×10^7^ parasites) were mixed with either *ΔACO* or *ΔIDH* parasites (≈5×10^7^ parasites) at 1:1 ratio respectively. Mixed cultures were harvested every 5 days over a course of 30 days and genomic DNA was isolated for PCR. Primer sets that specifically amplify the *hDHFR* gene (568 bp) and *PfGAPDH* gene (150 bp) are shown in S1 Table. The *hDHFR* PCR product indicated the presence and qualitative level of knockout parasites in the mixed culture, and the *PfGAPDH* PCR product was used as an internal control representing the presence and qualitative level of the total (wild type plus knockout) parasites present.

### Growth inhibition assay

Growth inhibition of WT and knockout parasites by the antimalarial compounds artemisinin, atovaquone and DSM-1 were measured by the [^3^H]-hypoxanthine incorporation assay in 96-well plates [34]. Briefly, parasites (predominantly ring-stage) at 1% parasitemia and 2% hematocrit were treated with each antimalarial compound at serial dilution concentrations ranging from 6000 nM to 0.1 nM for 24 h. Each well was pulsed with 0.5 μCi of [^3^H]-hypoxanthine and incubated for another 24 h. Parasites were then lysed by freezing and thawing. Nucleic acids were collected on filters with a cell harvester (PerkinElmer Life Sciences). After addition of MicroScint O (PerkinElmer Life Sciences), incorporation of [^3^H]-hypoxanthine was quantified using a TopCount scintillation counter (PerkinElmer Life Sciences).

### Gametocyte induction and exflagellation assay

For gametocyte cultures, asynchronous asexual stage NF54 cultures (WT and knockout lines) at 5-10% parasitemia were split with fresh RBCs and seeded at 0.5% parasitemia at 2.5% hematocrit in a 50 ml volume on day 0. The standard RPMI-1640 medium was supplemented with 5% O^+^ human serum (Key Biologics, LLC (Memphis, TN)) and 0.25% AlbuMAX. The gametocyte cultures were fed daily without being split for up to 20 days. For antioxidant treatments, N-acetylcysteine (NAC) at 1 mM, 5 mM or 10 mM and MitoQ at 100 nM were added into the medium daily from day 2 to day 20. For citrate treatment, the medium was supplemented with 500 μM sodium citrate dehydrate (Sigma) and used to feed the gametocyte cultures from day 2 to day 20. Gametocytemia was monitored daily by counting the number of gametocytes of all stages in 1,000 erythrocytes in Giemsa-stained thin blood smears. On days 14-20, for each gametocyte culture, the total number of erythrocytes was determined by a hemocytometer. The total number of gametocytes was then calculated by multiplying the number of erythrocytes by the gametocytemia. On each of these days, 1 ml of the cultures was collected, centrifuged briefly and resuspended in 200 μl exflagellation-inducing medium containing 25% human serum and 50 μM xanthurenic acid [35] at pH 8.4. After 10 min incubation at room temperature, 10 μl of the cell suspension was pipetted into a hemocytometer. The number of the exflagellation centers was counted using a hemocytometer under a standard optical microscope (X20 objective) over the next 10 min. The total number of exflagellation centers in each culture is calculated. Then the exflagellation percentage is calculated by dividing the total number of exflagellation centers by the total number of gametocytes in each culture.

### Gametocyte Percoll enrichment and mitochondrial staining

Gametocyte enrichment by Percoll was carried out using the method described in [36]. Briefly, on day 9-10 of the gametocyte induction, the cultures were harvested, centrifuged and resuspended to 50% hematocrit in complete RPMI-1640 medium. The cell suspension (2 ml) was then centrifuged through a Percoll step gradient in a 15 ml tube (from bottom to top: 2 ml of 80%, 2 ml of 65%, 2 ml of 50% and 2 ml of 35% Percoll solutions). The gametocyte-containing fraction at the 35%-50% interphase of the gradient was collected, washed three times with medium and resuspended with complete RPMI-1640 medium. Enriched gametocytes were then stained with 60 nM Mitotracker^®^ Red CMXRos (ThermoFisher Scientific) for 10 min at 37 °C and washed three times with medium. The gametocyte pellet was resuspended with an equal volume of complete RPMI-1640 medium and 3 μl of the suspension was pipetted onto a glass slide with a coverslip added on top. The gametocytes were imaged at room temperature using a 60X oil lens on a Nikon Eclipse T_i_-E fluorescence microscope. The images of bright field and DsRed channels were captured by a Nikon DS-Fi1c camera and processed with the NIS-Elements imaging software. The imaging procedure from loading the slide to image capture was finished within 15 min for each slide.

## RESULTS

### Knockout of *IDH* Gene in *P. falciparum*

We carried out the genetic disruption of *IDH* (PF3D7_1345700) in the gametocyte-competent NF54 strain of *P. falciparum* via double cross-over homologous recombination. As initial attempts using previous standard double crossover methods [25, 31, 33] were unsuccessful, we employed CRISPR-Cas9 genome editing, as adapted for use in *P. falciparum* [37], to achieve the knockout of *IDH* in NF54. Based on the criteria for potent single-guide RNAs (sgRNAs) described in [38], three independent *IDH*-specific sgRNA sequences were selected (S1 Table) and individually cloned into the pAIO plasmid [32], which carries the expression cassette for gRNA and the gene encoding the endonuclease Cas9. To provide template for homology-driven DNA repair, 5’ and 3’ HR DNA fragments from the *IDH* gene (S1 Fig panel A) were cloned into the pCC1 vector, as described in [25]. For transfection, we linearized the template pCC1-PfIDH vector with HincII and co-transfected NF54 ring-stage parasites with the linearized vector and all three sgRNA-Cas9 vectors. As shown in S1 Fig, we successfully generated *IDH* knockout (*ΔIDH*) parasites, in which a methotrexate-resistant *hDHFR* cassette replaced the target region of the *IDH* genomic locus. Diagnostic PCR using primers outside of the 5’HR and 3’HR (S1 Table) showed the expected product size, confirming the genotype of the *IDH* knockout, with no detectable wildtype band present (S1 Fig panel B).

### Phenotypic characterization of *ΔIDH* parasites in the asexual blood stage

To examine the phenotype of *ΔIDH* parasites, we measured the growth rate of tightly synchronized asexual parasites by monitoring parasitemia daily over a course of 8 days (4 intraerythrocytic cycles) by flow cytometry. *ΔIDH* parasites displayed a reduced growth rate trend compared to wild type (WT) NF54 parasites (Fig 1A). The *ΔIDH* parasites progressed through all the intraerythrocytic stages (ring, trophozoite and schizont stages) with apparently normal, healthy morphology, indicating that IDH is not essential in the asexual blood stage of *P. falciparum*. This is consistent with the previous finding that the *IDH* gene can be deleted in the gametocyte-incompetent D10 strain [25]. To corroborate the possible slower growth rate of *ΔIDH* parasites, we assessed long-term relative fitness of the parasites using a growth competition approach similar to previously reported in vitro competition experiments [39]. Equal numbers of WT and *ΔIDH* parasites (≈5×10^7^ parasites) were mixed in the same culture and allowed to compete for a growth period of 30 days. DNA samples were collected every 5 days from the culture, and the relative amount of KO parasite DNA present was monitored by PCR amplification of the *hDHFR* gene, which is only present in the *ΔIDH* parasites. A housekeeping gene, *P. falciparum* glyceraldehyde 3-phosphate dehydrogenase (*GAPDH*), was used as a total parasite DNA control in separate PCR reactions. As shown in Fig 1B, the intensity of the *hDHFR* PCR product decreased drastically after day 10 and became undetectable after day 20, while the *PfGAPDH* PCR product intensity remained essentially constant over the course of 30 days. These data revealed that there was a continuous loss of *ΔIDH* parasites in the mixed culture, indicating a significant fitness cost associated with the disruption of the *IDH* gene. Overall, we showed that, while *IDH* is not required for the viability of the asexual blood-stage *P. falciparum* parasite, its disruption in the NF54 line resulted in a fitness cost that reduced the growth rate.

### Sensitivity of *ΔIDH* and *ΔACO* parasites to antimalarial drugs that potentially target mitochondrial functions

The exact mechanism of action of artemisinin remains unsettled and may be multimodal [40, 41]. Based on findings in yeast and *P. falciparum* parasites [42, 43], it has been suggested that artemisinin induces reactive oxygen species (ROS) locally within the mitochondrion, which may be one mode of action. Since ACO and IDH are responsible for producing mitochondrial NADPH, which is thought to be required for maintenance of redox balance, as well as acting as a cofactor in biosynthetic reactions, we tested the possibility that knockout of *ACO* or *IDH* might hypersensitize the parasites to artemisinin. Growth inhibition assays, however, showed that the WT and KO parasites had similar susceptibility to artemisinin (Table 1). It has been shown that inhibition of the mtETC at complex III by atovaquone abolishes carbon flux through the TCA cycle [25]. We wondered whether asexual parasites that had adapted to knockout of *ACO* or *IDH* could have different responses to atovaquone. However, the EC_50_ values of atovaquone (shown as 95% confidence intervals) against the WT and KO lines are very similar (Table 1). In asexual blood-stage malaria parasites, the critical function of the mtETC was shown to be the regeneration of oxidized ubiquinone to serve as the electron acceptor for DHODH, an essential enzyme in the pyrimidine biosynthesis pathway [44]. We thus compared the sensitivity of the WT and KO parasites to the DHODH inhibitor DSM-1. Our results revealed no significant difference in the EC_50_ values of DSM-1 against WT or KO parasites, indicating that knockout of *ACO* or *IDH* does not affect the activity of DHODH. Altogether, the results of drug profiling assays indicate that KO of *IDH* or *ACO* did not significantly alter the sensitivity of the parasites to antimalarials that target the mitochondrion.

**Table 1.**
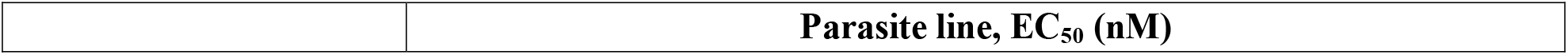
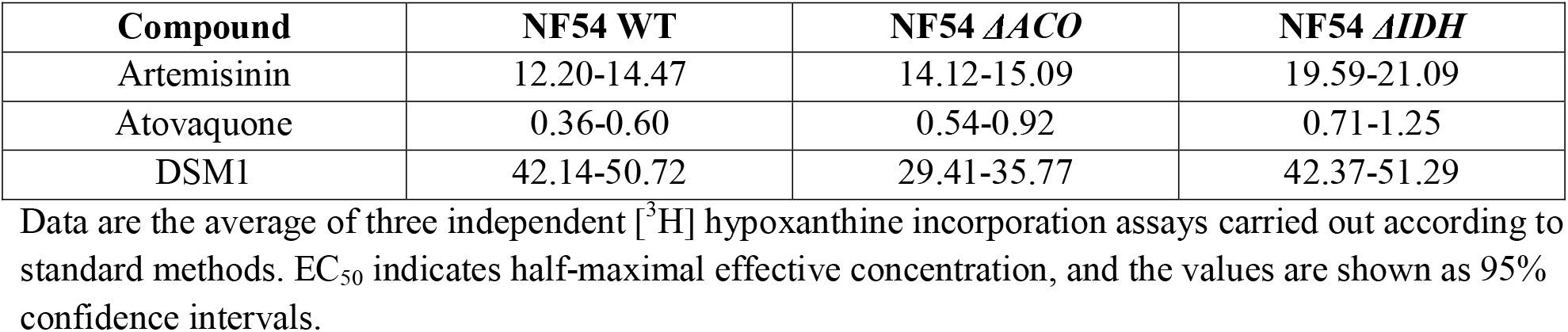
EC_50_ values of selected antimalarials vs. *ΔIDH* and *ΔACO* lines.

|  | Parasite line, EC<sub>50</sub> (nM) |
| --- | --- |

| Compound | NF54 WT | NF54 *ΔACO* | NF54 *ΔIDH* |
| --- | --- | --- | --- |
| Artemisinin | 12.20-14.47 | 14.12-15.09 | 19.59-21.09 |
| Atovaquone | 0.36-0.60 | 0.54-0.92 | 0.71-1.25 |
| DSM1 | 42.14-50.72 | 29.41-35.77 | 42.37-51.29 |

Data are the average of three independent [<sup>3</sup>H] hypoxanthine incorporation assays carried out according to standard methods. EC<sub>50</sub> indicates half-maximal effective concentration, and the values are shown as 95% confidence intervals.

### Phenotypic characterization of *ΔIDH* parasites in the gametocyte stages

To evaluate the effect of the knockout of *IDH* on gametocyte development, we induced gametocyte formation in NF54 WT and *ΔIDH* asexual parasites using standard methods (see “Materials and methods”). Maturation of *P. falciparum* gametocytes occurs over 10-14 days and is conventionally divided into five morphologically distinct stages (reviewed in [45]). While stage I resembles the asexual trophozoite stage, stage II starts to elongate and is quite distinguishable from asexual parasites. The parasites elongate further and become D-shaped at stage III. Stage IV gametocytes continue to elongate, developing pointed ends. Finally, mature stage V gametocytes acquire their typical crescent shapes with rounded ends. Normal morphologies of stage II-V WT gametocytes are shown in representative images of Giemsa-stained thin blood smears (Fig 2A). In contrast to normal, healthy WT gametocytes, *ΔIDH* gametocytes exhibited degenerated morphologies starting at stage III, indicated by reduced Giemsa staining intensity, which suggests loss or marked condensation of nucleic acid-containing structures (Fig 2A). Some stage III gametocytes progressed a little further to become stage IV-like parasites; however, none of them showed normal morphologies (Fig 2A). Most importantly, no mature stage V gametocytes were observed in *ΔIDH* cultures. Quantification of gametocytemia (percentage of gametocytes in the total number of erythrocytes) showed that the NF54 WT line produced a greater number of gametocytes than the *ΔIDH* line consistently from day 7 to day 18 post gametocyte induction (Fig 2B). The diminished gametocytemia of the *IDH* KO line appears to be largely due to the arrest of gametocyte development at stage III/IV, followed by parasite death and disintegration.

**Fig 2.**
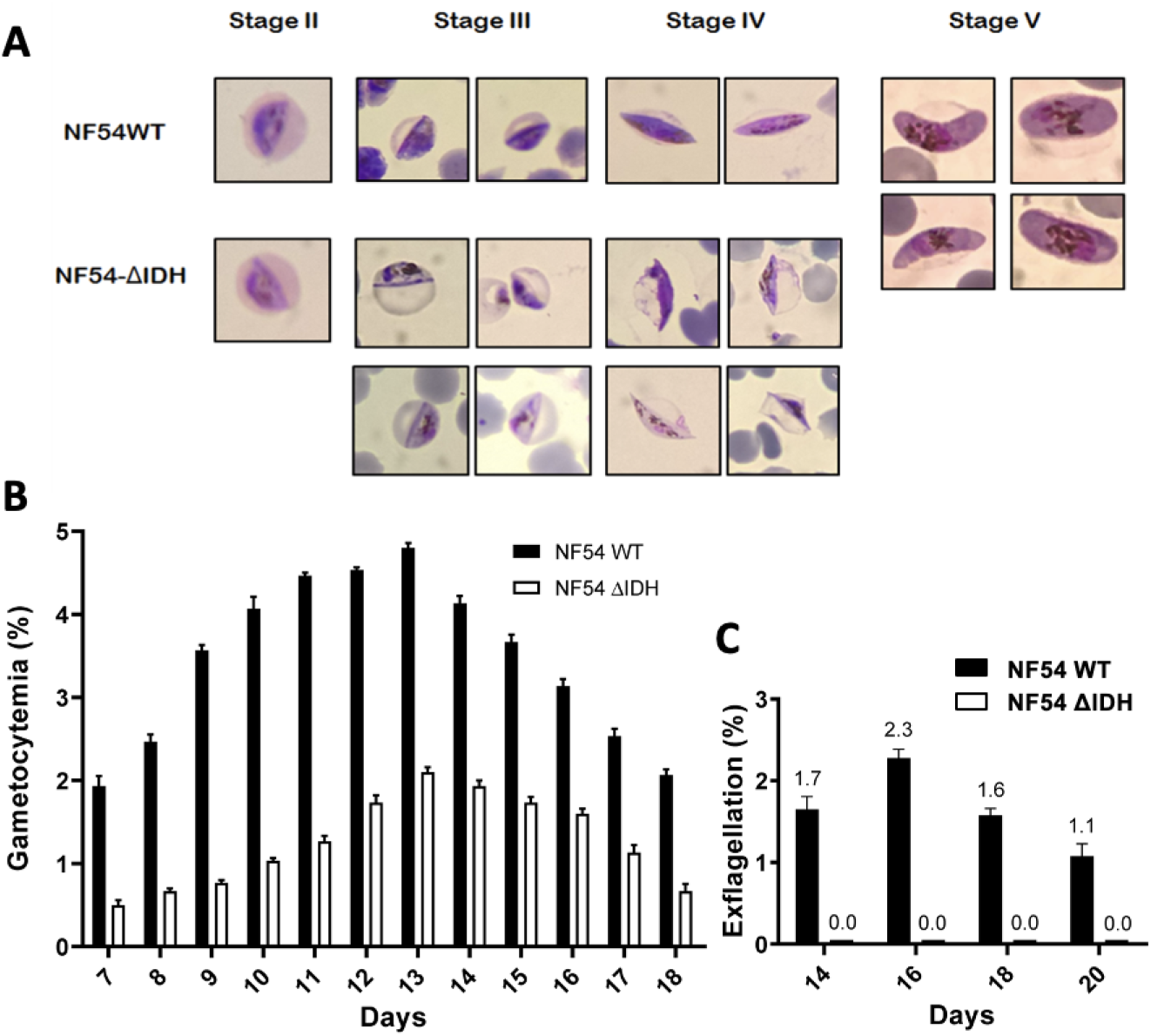
Phenotypic characterization of *ΔIDH* parasites in the gametocyte stage. (A) Representative Giemsa-stained images of NF54 WT (top row) and NF54 *ΔIDH* (bottom row) gametocytes at various development stages (II-V). (B) Gametocytemia of NF54 *ΔIDH* parasites is consistently lower than that of NF54 WT on various days during the gametocyte development. Gametocytemia is the percentage of gametocytes of all stages determined by counting 1000 RBCs. (C) NF54 *ΔIDH* line is defective in male gamete formation. Exflagellation percentage is the fraction of exflagellating gametocytes determined from 1000 gametocytes of all stages. Results are averaged from three independent experiments (error bars show the SEM, standard error of the mean.).

Since no stage V gametocytes were observed in *ΔIDH* parasites, formation of gametes was not expected. To confirm this, we attempted to induce exflagellation in the cultures from days 14 to 20. Exflagellation is the process by which mature stage V male gametocytes undergo gamete formation, which involves three rounds of DNA replication within 8-10 minutes and rapid protrusion of eight flagellum-like microgametes. Exflagellation is a hallmark of healthy mature stage V male gametocytes. The exflagellation process can be induced in vitro by a temperature drop [46, 47], a pH increase from 7.4 to 8.0-8.2 [46, 48] and a low concentration of xanthurenic acid [35]. The highly motile male gametes tend to bind surrounding erythrocytes and form moving cell clusters (exflagellation centers) that are visible and countable under a light microscope. We assessed exflagellation of NF54 WT and *ΔIDH* gametocytes by counting exflagellation centers. The *ΔIDH* line did not form any exflagellation centers, while the percentage of exflagellation centers over the total number of gametocytes in the WT line varied from 1-2.5% (Fig 2C). Thus, ablation of IDH resulted in abortive gametocyte development with abnormal late-stage morphologies, similar to the results reported for *ΔACO* parasites [25]. These results suggest that *IDH* is essential for gametocytogenesis in *P. falciparum* parasites.

## Mitochondrial morphology in *ΔACO* and *ΔIDH* gametocytes

Since IDH and ACO are both mitochondrion-localized TCA cycle enzymes, we examined the mitochondrial morphologies of the KO lines by Mitotracker staining in live gametocytes enriched by Percoll (see “Materials and methods”). Despite the indistinguishable mitochondrial morphology of asexual blood-stage NF54 WT and KO parasites, we observed drastically different mitochondrial structures in KO gametocytes (Fig 3A). In the WT stage III/IV gametocytes, the single mitochondrion had a long filamentous structure, which is consistent with previous observations in gametocytes with GFP-labeled mitochondria [22]. Interestingly, the mitochondria in *ΔACO* and *ΔIDH* gametocytes appeared to be fragmented, as indicated by the punctate staining patterns (Fig 3A). Approximately 100 gametocytes from each parasite line were categorized based on their mitochondrial morphology as either filamentous or fragmented. Quantification showed that more than 95% of the WT stage III/IV gametocytes had long filamentous mitochondria while 96% of *ΔACO* gametocytes and 99% of the *ΔIDH* gametocytes had fragmented mitochondria (Fig 3B). As WT gametocytes have been observed to contain a single mitochondrion [49], which elongates and branches during maturation, the fragmented morphology seen in the gametocytes of the KO lines appears to indicate some form of mitochondrial dysfunction and/or damage.

**Fig 3.**
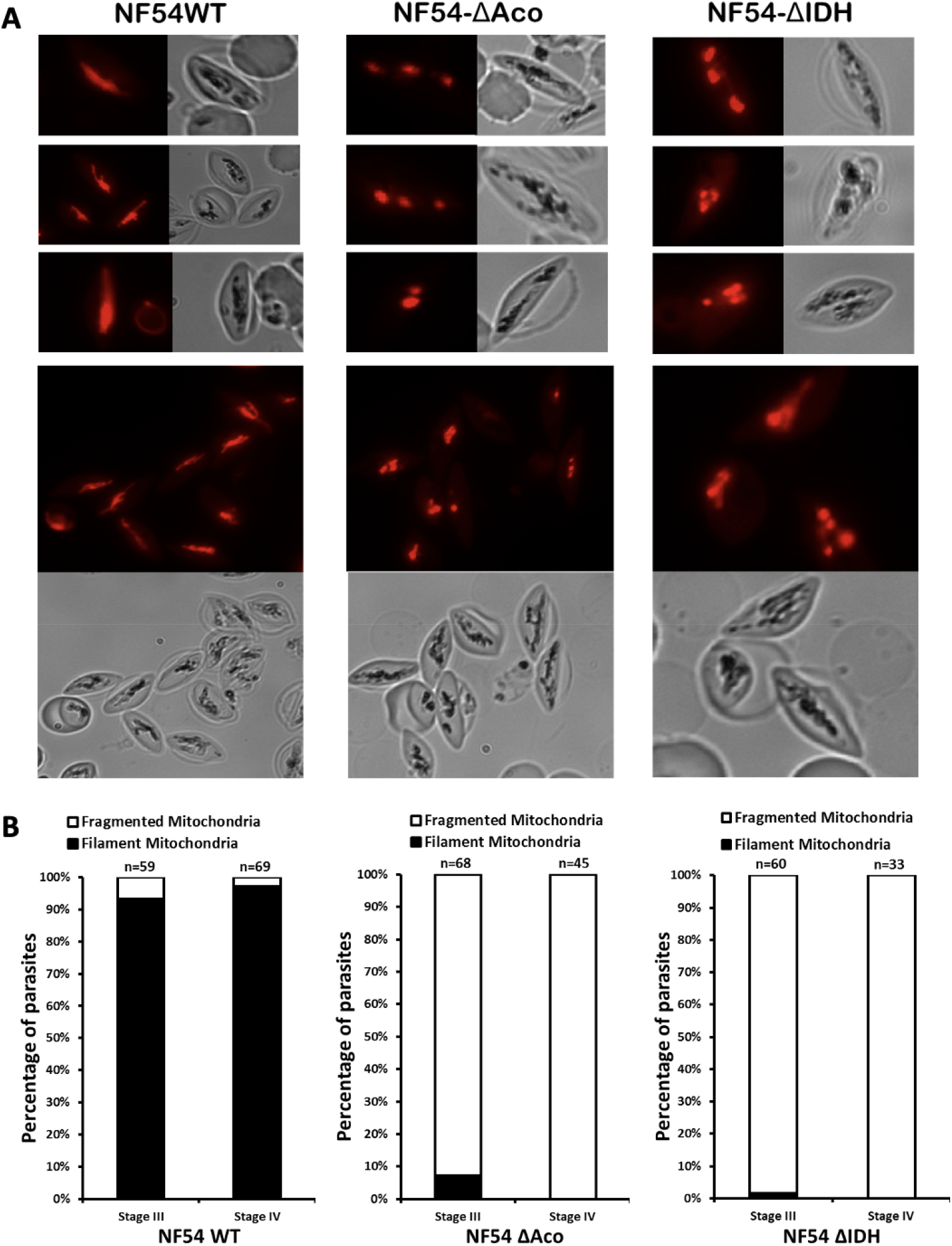
*ΔACO* and *ΔIDH* gametocytes exhibit fragmented mitochondrial morphology. (A) Representative mitotracker-stained images of NF54 WT (left), NF54 *ΔACO* (middle) and NF54 *ΔIDH* (right) live gametocytes on day 9-10 of the gametocyte development. (B) Quantification of the numbers of gametocytes with filamentous or fragmented mitochondria, respectively, in mitotracker-stained NF54 WT (left), NF54 *ΔACO* (middle) and NF54 *ΔIDH* (right) live gametocytes. n is the total number of the gametocytes counted in each case.

### Effect of antioxidant treatments

The above results with *ΔACO* and *ΔIDH* parasites have revealed a crucial role of the short TCA cycle segment from citrate to α-ketoglutarate in gametocyte development. We previously showed that knockout of *KDH* abolished the major flux through theTCA cycle but did not lead to any observable defects in gametocyte development [25], suggesting that a complete TCA cycle is not required for gametocytogenesis. Thus, we reasoned that the individual IDH and ACO enzymatic activities (or the very short ACO–IDH pathway), rather than their involvement in the overall TCA cycle, are crucial for gametocytogenesis. There appear to be two main consequences of eliminating ACO or IDH: (1) the accumulation of upstream citrate and (2) the ablation of mitochondrial NADPH produced by IDH. Therefore, we tested both possibilities in the context of gametocyte development.

First, we attempted to rescue the defective phenotype in *ΔACO* and *ΔIDH* gametocytes with antioxidant treatments, since the lack of mitochondrial NADPH could disrupt the local redox balance. NADPH is not membrane permeable; therefore, it cannot be simply added into the parasite medium. We first used NAC, a water-soluble antioxidant that is widely used *in vivo* and *in vitro* and provides a source of cysteine, which is an important precursor for biosynthesis of the primary intracellular antioxidant and detoxification agent glutathione [50-53]. We found that 1 mM and 5 mM NAC slightly increased the overall gametocytemia of *ΔACO* gametocytes, but the level was still significantly lower than the WT groups (Fig 4A). Nevertheless, NAC, up to 10 mM, did not restore any gamete formation in *ΔACO* gametocytes, indicating that there were no mature stage V gametocytes formed (S2 Table tab A). Next, we utilized MitoQ, a mitochondria-targeting antioxidant. This compound contains a lipophilic and positively charged triphenylphosphonium (TPP) group, which allows it to pass through membranes and accumulate near the negatively charged inner leaflet of the mitochondrial membrane. The reduced form of the ubiquinone head group of MitoQ can detoxify free radicals or ROS and is then recycled back to the reduced form by the action of electron transport chain complex II and possibly other quinone-dependent dehydrogenases [54, 55]. However, we did not observe any effect of 100 nM MitoQ on the overall gametocytemia or gamete formation by *ΔACO* or *ΔIDH* parasites compared to the groups treated with vehicle and non-antioxidant TPP compound (Fig 4B and 4C; S2 Table tabs B and C). Thus, the antioxidants tested were ineffective in overcoming the deleterious effects of the loss of IDH or ACO on gametocyte development. However, it is possible that externally provided NAC is not efficiently taken up by the parasites, and MitoQ can potentially exhibit pro-oxidant activity (see Discussion), so these results may not prove definitive.

**Fig 4.**
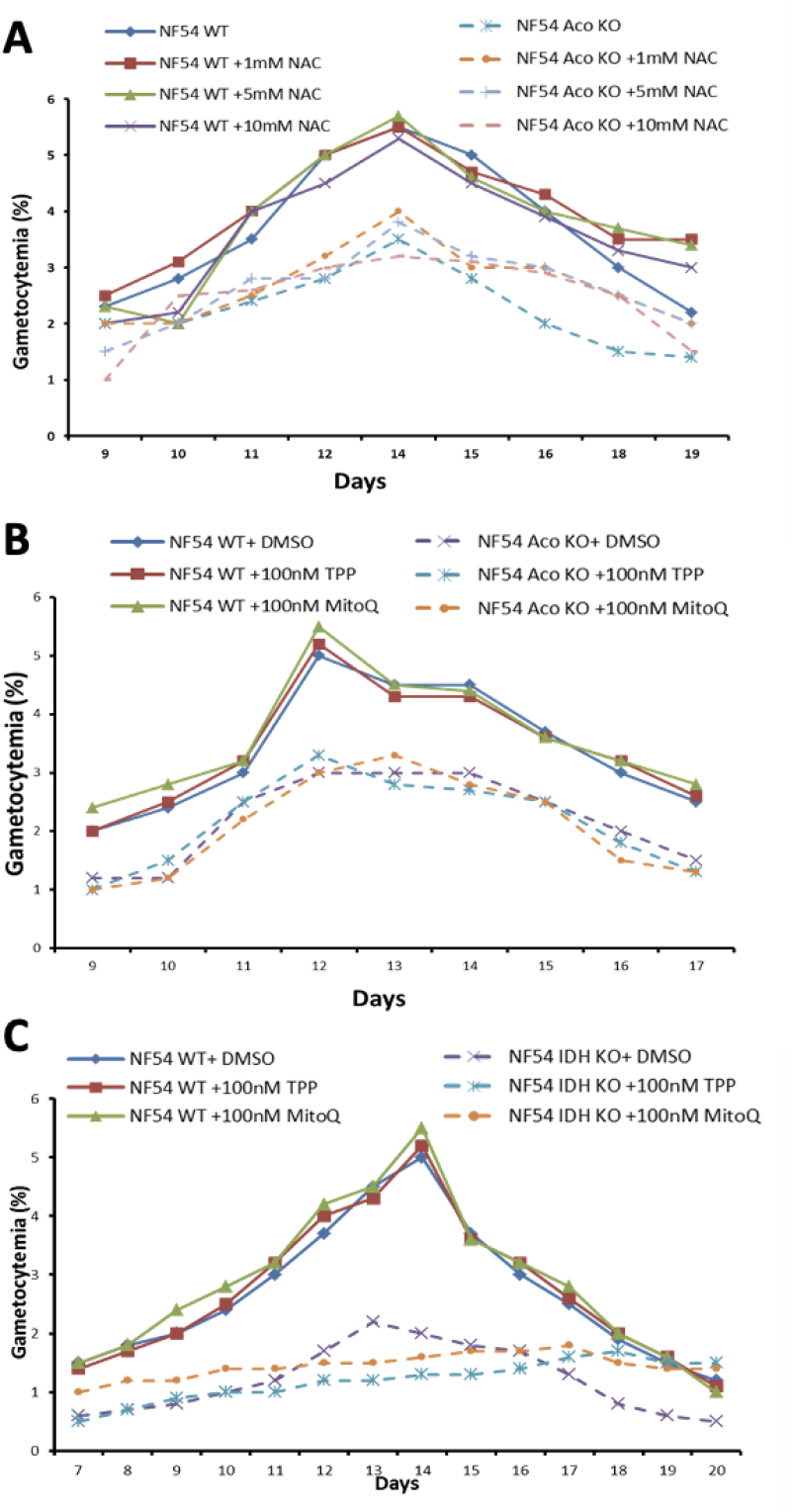
Antioxidants failed to rescue the defective phenotype in *ΔACO* and *ΔIDH* gametocytes. (A) Gametocytemia of NF54 *ΔACO* parasites is not restored by NAC at 1 mM, 5 mM or 10 mM compared to treated and non-treated WT gametocytes. (B) Gametocytemia of NF54 *ΔACO* parasites relative to WT parasites is not restored by 100 nM MitoQ. Triphenylphosphonium (TPP) is used as a mitochondrion-targeted non-antioxidant control for MitoQ. (C) Gametocytemia of NF54 *ΔIDH* parasites is not restored by 100 nM MitoQ. Results are averaged from three independent experiments (error bars show the SEM, standard error of the mean).

### Effect of citrate treatment on gametocyte development and gamete formation

Another consequence of inhibiting the oxidative arm of the TCA cycle is the accumulation of upstream metabolites isocitrate/citrate. Toxicity of excessive amounts of citrate has been observed in yeast, potentially due to disruption of iron homeostasis [56]. Metabolite analysis by gas chromatography-mass spectrometry (GC-MS) showed that chemical inhibition of aconitase by the NaFAc led to a 17-fold increase of citrate in gametocytes [26]. Furthermore, an increased level of citrate/isocitrate (detected by LC-MS as the same metabolite due to their identical masses) was excreted into the growth medium by the *ΔIDH* asexual parasites (D10 strain) compared to the WT parental line [25]. This suggests the permeability/transport of citrate across both the parasite plasma membrane and mitochondrial membranes. Therefore, we investigated the possibility of citrate toxicity by directly adding a relatively high concentration of citrate into the culture medium of WT gametocytes. Gametocyte cultures of NF54 WT parasites were treated with 500 μM citrate throughout the 20-day gametocyte assay, and gametocytemia and gamete formation were monitored. Citrate treatment did not have any significant effect on gametocyte development or gamete formation in the NF54 WT parasites (Fig 5), suggesting that the buildup of citrate is likely not the cause of the defective phenotypes that we observed in the NF54 *ΔACO* or *ΔIDH* gametocytes.

**Fig 5.**
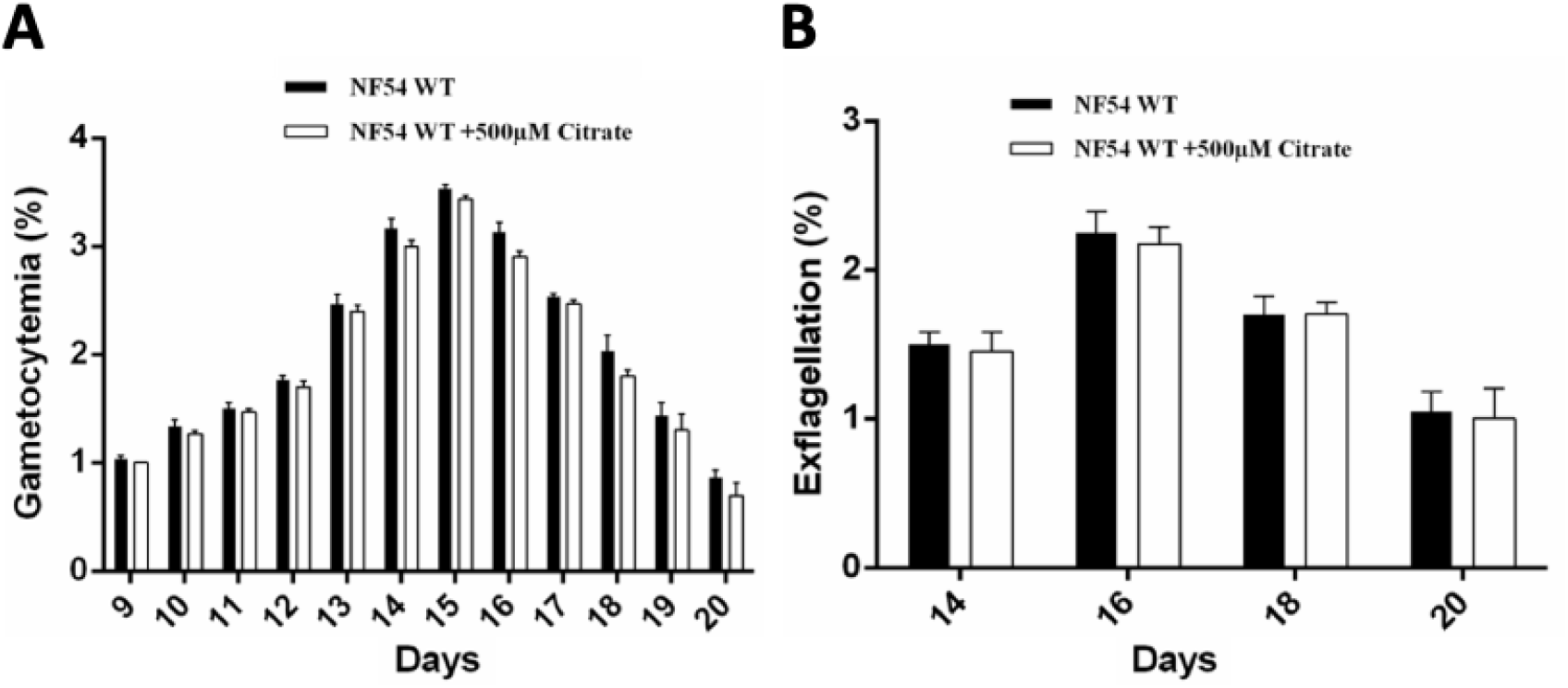
Citrate treatment does not affect gametocyte development or gamete formation of NF54 WT parasites. (A) Day 9-20 gametocytemia of NF54 WT parasites that were untreated or treated with 500 μM citrate. (B) Exflagellation percentages of NF54 WT gametocytes that were untreated or treated with 500μM citrate on day 14, 16, 18 and 20 of the gametocyte induction assay. Results were averaged from three independent experiments. (Error bars show the SEM, standard error of the mean).

## Discussion

Sexual stage development is an important component of the complex malaria parasite life cycle that allows the parasites to leave their vertebrate hosts, undergo sexual reproduction in the mosquito vector and subsequently to infect new hosts. From a therapeutic standpoint, understanding the biology of the sexual stages is a key to develop drug leads that can block malaria transmission. A crucial role of the parasite mitochondrion in gametocyte development has been indicated for many years by various studies on cellular, genetic, and biochemical levels [20-23, 25, 26]. However, our knowledge on which aspects of mitochondrial functionality are involved is still limited. Previous observations revealed the functional importance of aconitase in gametocyte development [25]. In the current study, we investigated the essentiality of another TCA enzyme–IDH. Altogether, we established the specific necessity of a partial TCA cycle during gametocytogenesis, i.e., from citrate to α-ketoglutarate catalyzed by ACO and IDH.

Compared to asexual blood-stage parasites, *P. falciparum* gametocytes are quite distinct in terms of their morphologies [20, 22, 45], gene expression profiles [23, 57], cellular development and metabolism (reviewed in [58]). Our study provides an example in which the functionality of a particular gene is critical for sexual development of the parasite, while the same gene is dispensable during the asexual blood stage. While *IDH* can be genetically disrupted with some fitness cost to the asexual blood stage (Fig 1), the sexual differentiation and gamete formation of the *ΔIDH* parasites were severely impaired (Fig 2) The necessity of *IDH* for gametocytogenesis justifies the evolutionary maintenance of the gene by *P. falciparum* because parasites without the ability to develop gametocytes will become extinct. This reinforces the finding that a complete TCA cycle is required in the insect stages of the parasite [25]. Interestingly, *ΔIDH* parasites showed a fitness cost during asexual replication, suggesting that IDH deficiency might cause a degree of damage, but the 48-hour asexual intraerythrocytic cycle is insufficient to manifest lethal consequences. However, during the gametocyte development, which typically lasts for 10-14 days, the damage may accumulate beyond the threshold of tolerance in the *ΔIDH* parasites. Our results once again showcased that mitochondrial functions become more important in the sexual development of the parasites compared to their asexual replication.

We observed the mitochondria in *ΔACO* and *ΔIDH* gametocytes with fragmented morphologies compared to the long filament structures in the WT mitochondria. The punctate staining of the mitochondria of *ΔACO* and *ΔIDH* gametocytes is likely the result of fragmentation of the single mitochondrion, rather than remnants of multiple mitochondria within the same gametocyte. Live cell imaging of *P. falciparum* gametocytes with GFP-labeled mitochondria previously revealed the presence of a single mitochondrion in each gametocyte that undergoes a large expansion to form branching structures as the gametocyte matures [22], leading Noriko Okamoto *et al*. [22] to remark that the highly lobed mitochondrion might appear as multiple separated organelles in 2D images, as previously described in other studies [20]. The Mitotracker staining patterns we observed in *ΔACO* and *ΔIDH* gametocytes, however, indicated that the single mitochondrion underwent fragmentation in these mutant parasites. The degree of dysfunction requires further study, but interestingly, these fragments maintained a membrane potential since they were still stained with Mitotracker, which only accumulates in negatively charged compartments.

Malaria parasites face oxidative threats throughout their life cycles. Upon ingestion into the mosquito midgut, *P. falciparum* parasites are exposed to high levels of ROS induced by the mosquito immune system [59]. In the blood circulation of a malaria-infected human host, monocytes produce pro-inflammatory cytokines and hydroxyl radicals as the first-line defense mechanism against the parasites [60]. More importantly, during the blood stage, the parasites need to cope with several major sources of ROS: H_2_O_2_ and hydroxyl and other free radicals generated from free heme released from hemoglobin digestion within the digestive vacuole and from hemichromes (oxidized hemoglobin containing reactive iron) formed in the infected host RBC [61-64]. In addition, the mtETC can leak electrons to oxygen, generating superoxide anions [65]. Therefore, a functional redox-balancing system is crucial for the survival of the parasites. In *P. falciparum*, a cytosolic glutathione and thioredoxin system is responsible for maintaining the redox homeostasis [28]. The redox system in the parasite mitochondria has not been fully characterized, but there are numerous lines of evidence for the presence of a functional glutathione and thioredoxin system within the parasite mitochondria, including the presence of a glutaredoxin-like protein (GLP3) [66], a thioredoxin-like protein (TLP2) [67], a thioredoxin reductase isoform (TRXR) [67] and a peroxiredoxin (Trx-Px2) [68]. NADPH is required as an electron donor to power both glutathione and thioredoxin systems. Furthermore, NADPH also serves as an important cofactor for many NADPH-dependent enzymes. One example involves the ferredoxin which catalyzes sulfur reduction during the *de novo* biosynthesis of iron-sulfur (FeS) clusters (ISC) in the mitochondrion [69]. NADPH is required by ferredoxin reductase to regenerate the reduced form of ferredoxin [70]. Therefore, mitochondrial NADPH is likely critical for FeS cluster assembly, which is required for the function of the mtETC and many other processes in the cytoplasm as well as in the nucleus. IDH may be the only source for mitochondrial NADPH. The two isoforms of NADP^+^-dependent glutamate dehydrogenase (GDH) were considered to be another potential source for mitochondrial NADPH, as isoforms of GDH are found in the mitochondrion in plants and mammals, but the *P. falciparum* isoforms were shown to be localized in the parasite cytosol and apicoplast, rather than the mitochondrion [71, 72]. IDH was also shown to be upregulated at mRNA and protein levels under oxidative stress [27], thus suggesting a role of IDH in the maintenance of mitochondrial redox balance in the parasite.

It is challenging to assess the role of mitochondrial NADPH by measuring mitochondrial ROS levels in gametocytes. Direct assessment of mitochondrial ROS levels in the gametocytes is difficult because large amount of ROS is produced in the parasite food vacuole [73], which overwhelms the signal from the mitochondria. Recently, a genetically encoded glutathione biosensor has been utilized to measure the redox potential in the cytosol, apicoplast and mitochondrion of *P. falciparum* parasites [74, 75]. This approach relies on the redox-sensitive GFP protein (roGFP2), which undergoes an excitation shift from 408nm to 488nm resulting from the formation of an intramolecular disulfide bond in response to increased ROS level in the environment. The redox condition in the mitochondrial matrix was found to be highly reducing (−328 mV) and largely unaffected by treatment with the mtETC inhibitor atovaquone with or without proguanil. Future investigations utilizing the mitochondrion-targeted glutathione redox biosensor in *ΔACO* and *ΔIDH* lines could reveal whether lack of mitochondrial NADPH production results in a significant change of the matrix glutathione redox potential toward oxidation in asexual and gametocyte KO parasites. Antioxidants such as N-acetylcysteine (NAC) and MitoQ failed to rescue the defective phenotype in KO gametocytes (Fig 4). It is possible that the parasite mitochondrion is not readily accessible to NAC. It has been argued that the effect of exogenous ubiquinones, such as MitoQ, is difficult to interpret as they can exhibit pro-oxidant activity or act as electron carriers to facilitate respiration [76]. Specifically, MitoQ was able to autoxidize to generate superoxide [76]. This might explain our MitoQ results, if the antioxidant effect of MitoQ is neutralized by its own pro-oxidant action in the KO gametocytes. It is also possible that the potential non-redox roles of mitochondrial NADPH, such as its involvement in the FeS assembly pathway, could lead to detrimental effects on *ΔACO* and *ΔIDH* parasites in addition to any effects of increased ROS levels. This might explain our observation that antioxidant treatments were not able to rescue the defective phenotype in the KO gametocytes. Another consequence of the loss of aconitase or IDH is the accumulation of upstream citrate. Indeed, Ke *et al*. [25] showed that genetic knockout of *IDH* in D10 parasite line led to an increase of extracellular citrate concentration from 0.15 μM to 0.6 μM. Citrate toxicity could potentially cause impaired gametocytogenesis. However, addition of 500 μM citrate in the culture medium throughout gametocyte development did not affect gametocytogenesis or gamete formation in the NF54 WT parasites. These results suggest that the defective phenotype in the *ΔACO* and *ΔIDH* gametocytes is not likely due to increased citrate accumulation.

In conclusion, our study demonstrates that the TCA cycle enzyme IDH is essential for gametocyte development and gamete formation in *P. falciparum* malaria parasites and is dispensable during the asexual blood stage of the parasites, although its absence does exert a fitness cost to the parasite. While the exact molecular basis behind its criticality in gametocytes requires further study, specific inhibitors targeting *P. falciparum* IDH might be effective against the sexual stage of the parasites, thus blocking malaria transmission and preventing further infections.

## Acknowledgements

We thank the Malaria Research and Reference Reagent Resource of the National Institutes of Allergy and Infectious Diseases for the provision of various *P. falciparum* lines. We thank Dr. Christian Sell at Drexel University College of Medicine for providing the MitoQ compound. We also thank Drs. Josh R. Beck and Daniel E. Goldberg at Washington University School of Medicine for the gift of the pAIO plasmid. Gametocyte-competent NF54 WT line was kindly provided by Dr. Marcelo Jacobs Lorena from Johns Hopkins University.

## Supporting information

**S1 Table. Primers and oligonucleotides used in this study**.

**S2 Table. Antioxidants fail to rescue exflagellation of** *Δ****ACO* and** *Δ****IDH* gametocytes**.

**S1 Fig. Knockout of the IDH gene in *P. falciparum*. (**A) Schematic of *IDH* knockout by CRISPR-Cas9 genome editing. The 5’-homologous sequence (5’HR) is 924bp long (−584 to +339) and the 3’-homologous sequence (3’HR) is 988bp long (−321 to +666). The small arrows and vertical lines indicate primers used for diagnostic PCR in B. 2.9kb and 4.3kb are the sizes of the expected PCR products from WT and IDH KO parasites respectively. DSBs, double strand breaks. (B) Diagnostic PCR assessing the genotype of *ΔIDH* parasites using the primers depicted in A (see S1 Table).

**S2 Fig. Full images of cropped DNA gels shown in Fig 1B**. Following electrophoresis, agarose gels were stained with ethidium bromide and imaged in a GE Healthcare LAS 4000 ImageQuant instrument. (A) *hDHFR* PCR products; (B) *PfGAPDH* PCR products.

