## Supplementary figures and images for "Genetic disruption of isocitrate dehydrogenase arrests the full development of sexual stage parasites in *Plasmodium falciparum*"

### Supplementary Fig. 1

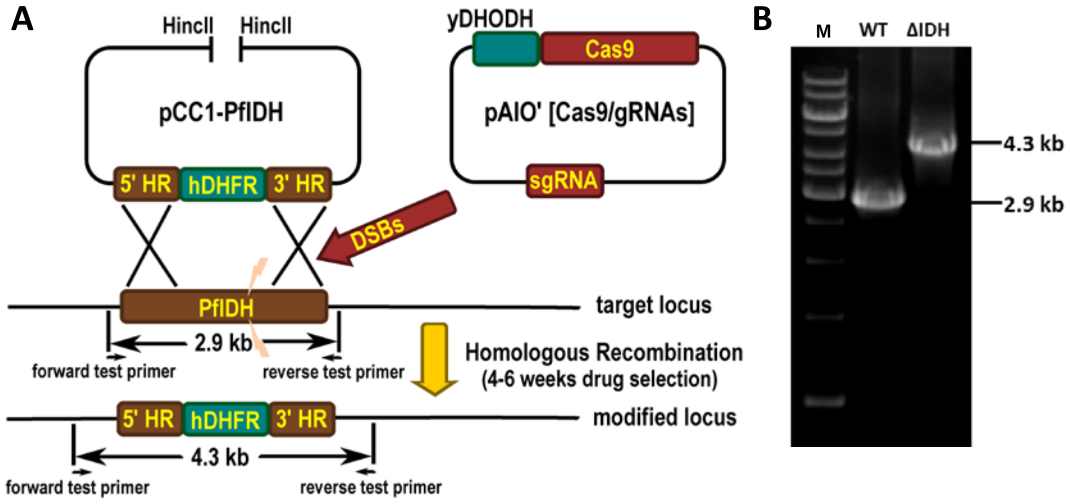
