## Supplementary Table 1 for "Genetic disruption of isocitrate dehydrogenase arrests the full development of sexual stage parasites in *Plasmodium falciparum*"

### S1 Table. Primers and oligonucleotides used in this study.

| Primer ID | Primer sequence | Purpose |
|---|---|---|
| PfIDH5fNcoI_F | GAccatggATCATCTTTATTTTTAGCGCACAC | For donor construct pCC1-PfIDH |
| PfIDH5fEcoI_R | CTgaattcCTTTAACTCTTGCAGCATCAGG | |
| PfIDH3fSpeI_F | GAactagtGCATGGACCAAAGGATTAGAAC | |
| PfIDH3fSacII_R | TAccgcggCAACTATGTTAAGGATAATACAGTATTG | |
| PfIDHKO_F | CTCCTGCTCATCAAAATGATAAATATG | Diagnostic PCR for IDH KO |
| PFIDHKO_R | TGGGAAAATTGTATTGCCTCA | |
| PfIDHgRNA1_OligoF | TAAGTATATAATATTGAAACACACGTAACACCGTCGTTTTAGAGCTAGAA | Cloning of IDH-specific gRNAs into Cas9 plasmid |
| PfIDHgRNA1_OligoR | TTCTAGCTCTAAAACGACGGTGTTACGTGTGTTTCAATATTATATACTTA | |
| PfIDHgRNA2_OligoF | TAAGTATATAATATTTGTGTTTCAGAAGCTGCCCAGTTTTAGAGCTAGAA | |
| PfIDHgRNA2_OligoR | TTCTAGCTCTAAAACTGGGCAGCTTCTGAAACACAAATATTATATACTTA | |
| PfIDHgRNA3_OligoF | TAAGTATATAATATTCTTGAGCAACAGCATCAGATGTTTTAGAGCTAGAA | |
| PfIDHgRNA3_OligoR | TTCTAGCTCTAAAACATCTGATGCTGTTGCTCAAGAATATTATATACTTA | |
| PfIDHgRNA1_N20 | GAAACACACGTAACACCGTC | gRNA sequences |
| PfIDHgRNA2_N20 | TGTGTTTCAGAAGCTGCCCA | |
| PfIDHgRNA3_N20 | CTTGAGCAACAGCATCAGAT | |
| gRNArevseq | TAGGAAATAATAAAAAAGCACC | gRNA sequencing |
| hDHFR_F | CAGGATCCATGCATGGTTCGCTAAAC | Diagnostic PCR for fitness cost assay |
| hDHFR_R | CGAAGCTTAATCATTCTTCTCATATAC | |
| PfGAPDH_F | TTTCCAATGCATCATGTACCA | |
| PfGAPDH_R | AGTCCTTACCACCCTTTGAT | |
